# SmartCodes: functionalized barcodes that enable targeted retrieval of clonal lineages from a heterogeneous population

**DOI:** 10.1101/352617

**Authors:** Clare Rebbeck, Florian Raths, Bassem Ben Cheik, Kenneth Gouin, Gregory J. Hannon, Simon R. V. Knott

**Author notes:** these authors contributed equally to this work (Clare Rebbeck, Florian Raths). to whom correspondence should be addressed (GJH, SRVK, CR).

## Abstract

Molecular barcoding has provided means to link genotype to phenotype, to individuate cells in single-cell analyses, to enable the tracking of evolving lineages, and to facilitate the analysis of complex mixtures containing phenotypically distinct lineages. To date, all existing approaches enable retrospective associations to be made between characteristics and the lineage harbouring them, but provide no path toward isolating or manipulating those lineages within the complex mixture. Here, we describe a strategy for creating functionalized barcodes that enable straightforward manipulation of lineages within complex populations of cells, either marking and retrieval of selected lineages, or modification of their phenotype within the population, including their elimination. These “SmartCodes” rely on a simple CRISPR-based, molecular barcode reader that can switch measurable, or selectable markers, on or off in a binary fashion. While this approach could have broad impact, we envision initial approaches to the study of tumour heterogeneity, focused on issues of tumour progression, metastasis, and drug resistance.

## Introduction

Mining the phenotypic heterogeneity of single cells present within cell populations is becoming increasingly important as a source of new insights into normal development and disease. In tumor-derived material and cell lines, sub clonal populations clearly differ in their properties, including the ability to contribute to different aspects of tumour initiation and progression and in their responses to therapy [1–3]. The use of genetic barcodes coupled with high throughput sequencing has facilitated the study of heterogeneity in many contexts [1, 4–6]; however, current methodologies are limited in that they provide only a catalogue of which cells exhibited particular behaviours. This usually comes without additional accompanying detail or any means to discern mechanisms underlying behavioural diversity. Importantly, genetic barcodes alone do not provide a means to isolate clonal populations from a heterogeneous mixture to enable further study.

Recent work has demonstrated the utility of dividing a population into its clonal components and studying their phenotypic diversity [4]. Genetic barcoding indicated the presence of clonal lineages within a mouse mammary tumour model, 4T1, that were proficient at forming circulating tumour cells, CTCs, with a reproducible subset of these being uniquely competent to form metastases at secondary sites. However, to probe the mechanistic basis of these phenotypes, clonal lineages had to be established by single-cell isolation and used to reconstitute a heterogeneity model from a limited initial diversity. Fortunately, those same phenotypes were observed in a collection of 23 clonal lines, enabling the use of molecular profiling to identify vascular mimicry as a driver of CTC forming potential and asparagine bioavailability as a driver of metastasis through its impacts on EMT [7, 8]

The need for manual isolation of clonal lineages as an intermediate step toward the study of heterogeneous populations raises a substantial barrier to the exploration of a plurality of models and the synthesis of insights derived from them. As an example, tumour cells vary in their sensitivity to therapeutic agents, ranging from sensitive to tolerant to resistant [9]. One can treat a population of tumour cells in vitro or in vivo and, for example, compare the expression profiles of resistant cells to the population as a whole. However, one cannot easily segregate these cell populations and study them independently, nor can one isolate those cells that are sensitive and compare their properties and expression patterns to resistant populations. Even if this scenario were successfully pursued by single-cell cloning with a given cell line, truly valuable insights would likely require the comparison of numerous models even within a single tumour subtype.

Here we present a CRISPR-based molecular barcode reader that when paired with functional barcodes (SmartCodes) enables the retrieval of selected clonal populations from heterogeneous mixtures. Like current barcoding methods, SmartCodes provide the ability to track the behaviour of clonal lineages by sequencing in complex samples. However, CRISPR-based activation of these functional barcodes also allows either selection for or selection against those clonal lineages in complex mixtures by using a cas9 deletion strategy to move an out of frame selection marker, in one case puromycin, into the coding frame. This enables enrichment of an sgRNA-targeted cell population that became puromycin resistant. Not only will this approach accelerate functional analysis of clonal lineages, it will also allow the isolation of even rare lineages with important phenotypic properties.

## Results and Discussion

### The SmartCode strategy

We considered that normally inert barcodes, could be functionalized simply by converting them into CRISPR target sites, in a way that interaction with Cas9 could elicit a predicted outcome. With respect to barcode design, this simply entails placing a known sequence of at least 20 nucleotides adjacent to a functional PAM site and appending that barcode to a genetic element that can confer the desired properties upon a cell. Cas9, programmed with an appropriate guide, would act as a molecular barcode reader and influence the activity of the genetic element. We term such functional barcodes, SmartCodes (Figure 1A).

**Figure 1.**
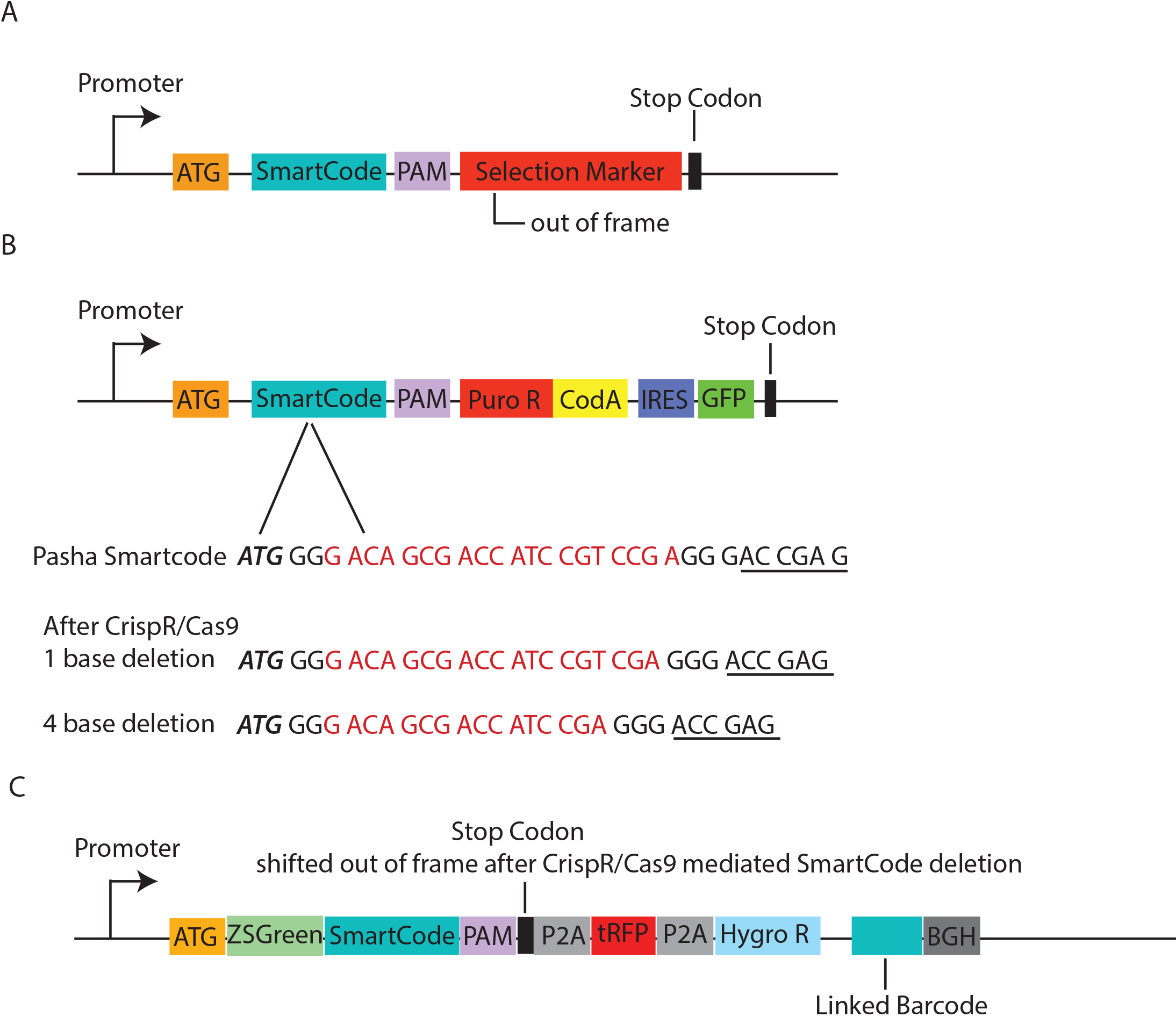
A) Basic design strategy for SmartCodes. B) SmartCode Vector with the Pasha sequence, and the dominant mutations after Cas9 targeting. Italics represents the start codon (ATG) and underlined are the first two codons of the puromycin-resistance gene. A spacer sequence (GG) was placed after the ATG and the PAM sequence (GGG) after the Pasha SmartCode (in red). C) Alternative vector design incorporating a conventional linked barcode adjacent to the polyA.

While there are many ways to achieve this goal, for example delivery of activators or repressors to a locus via a catalytically inactive Cas9, we chose to take advantage of the ability of Cas9 to cleave its targets and induce insertion or deletion (indel) mutations following repair by non-homologous end joining. By placing the genetic element in the appropriate context, its activity could be switched on or off in a binary fashion by shifting it into or out of a translational reading frame.

As one example, to purify a lineage of interest from a complex mixture, a SmartCode could be placed between a translational initiation site and a selectable maker, for example the puromycin-resistance gene, such that puro^r^ is out of frame and SmartCoded cells remain drug-sensitive. Action of Cas9 on the specifically targeted SmartCode would result in a subset of outcomes shifting the selectable marker into the correct frame, creating a puryomycin-resistant lineage that could be selected from a complex mixture by application of the drug. (Figure 2 outlines an example strategy for use of Smartcodes)

**Figure 2.**
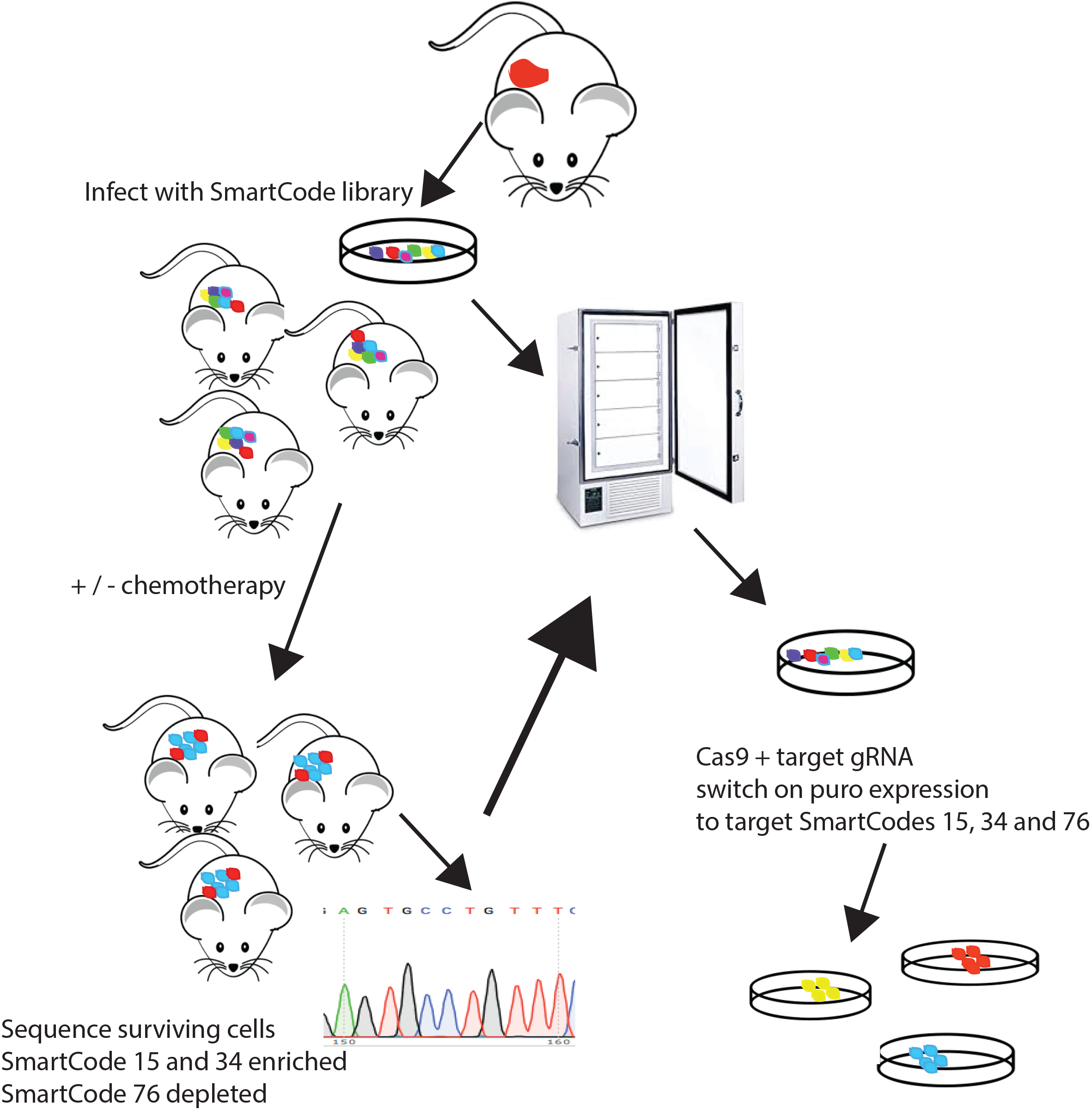
An example experiment where one can use Smartcodes to look for drug sensitive and drug resistance cells.

### Validation of the SmartCode approach

To validate our strategy, we devised a series of experimental simulations to test our ability to purify clonal lineages from mixtures of varying complexities. In our experience, and that of others, Cas9 targeting of a site often results in the generation of relatively small indels with each site having a set of reproducible, preferred outcomes [10]. We had previously characterized preferred outcomes for two target sites, one derived from GFP [11] and another from the human gene encoding the Pasha protein. Each had preferred outcomes (−1 or −4 deletion) that would shift a puromycin resistance marker designed in the +2 reading frame, into the +1 frame, consequently activating its expression.

These sites were used to create SmartCode selection cassettes in which the GFP (G) or Pasha (P) target sites were placed between an optimal initiator ATG and an out-of-frame resistance gene (Figure 1B). Selection cassettes were present downstream of a constitutive promoter in a retroviral vector that directs expression also of GFP, and derived viruses were used to infect 293T cells such that the overwhelming majority of cells had a single integrant. Infected cells were FACS sorted based upon the expression of the GFP marker to create populations in which most cells (85%) were infected. The two infected cell populations, harbouring GFP- or Pasha-derived SmartCodes, were then mixed at ratios of 1:500, 1:1000 and 1:10,000 to create populations that were tagged with a majority of GFP SmartCodes and a minority of Pasha SmartCodes.

In early trials, we noted that even without targeting the SmartCode with an appropriate guide RNA, we saw a substantial background of puromycin-resistant cells. As reverse transcription is an error-prone process, we hypothesized that frame shifts might be introduced into a small fraction of infected cells through this process, creating puro-resistant clones independently of any CRISRP-mediated event. In fact, deep sequencing of naive infected cell populations revealed mutations in SmartCode vectors that likely arose during virus production (not shown).

To address the issue of background resistance, we further modified our vector to place a negative selection marker in the same reading frame as the puromycin-resistance gene. Because it can be applied in many cellular contexts, we chose the cytosine deaminase gene (CodA) that converts 5-fluorocytosine to the toxic compound 5-fluorouracil. We used a specific CodA mutant, codAD314A [12], which shows higher activity and consequently increased toxicity in the presence of 5-FC. This dual selection cassette (Figure 1B) allows either positive or negative selection upon shift into the correct reading frame, and importantly could be used to eliminate cells where such a shift had occurred prior to exposure to the Cas9-based barcode reader.

To test SmartCode-based enrichment in our models of complex, heterogeneous populations, cells were first treated for 3 days with 5-FC to suppress unwanted background and then left to recover for a further 4 days. Populations were then infected with a virus encoding Cas9 and an sgRNA targeting the Pasha-derived site. After 7 days, the cells were treated with puromycin to select those in which the SmartCode has been targeted to activate the selectable marker. After further expansion of resistant cells (7 days), genomic DNA was extracted from the remaining cells and SmartCodes were amplified for sequencing.

Even without the use of 5-FC to suppress background, we saw a significant enrichment of the minority population (150 fold from a 1:1000 dilution). However, with background suppression, enrichment improved substantially. As an example, we could enrich a cell SmartCoded with a Pasha target site ~700 fold in a 1:1000 dilution of cells marked with a GFP SmartCode (Figure 3). The majority of mutations were a 1 or 4 base deletion (Figure 1B). These experiments demonstrated the ability to functionalize barcodes and to use these SmartCodes to enrich rare lineages from complex mixtures.

**Figure 3.**
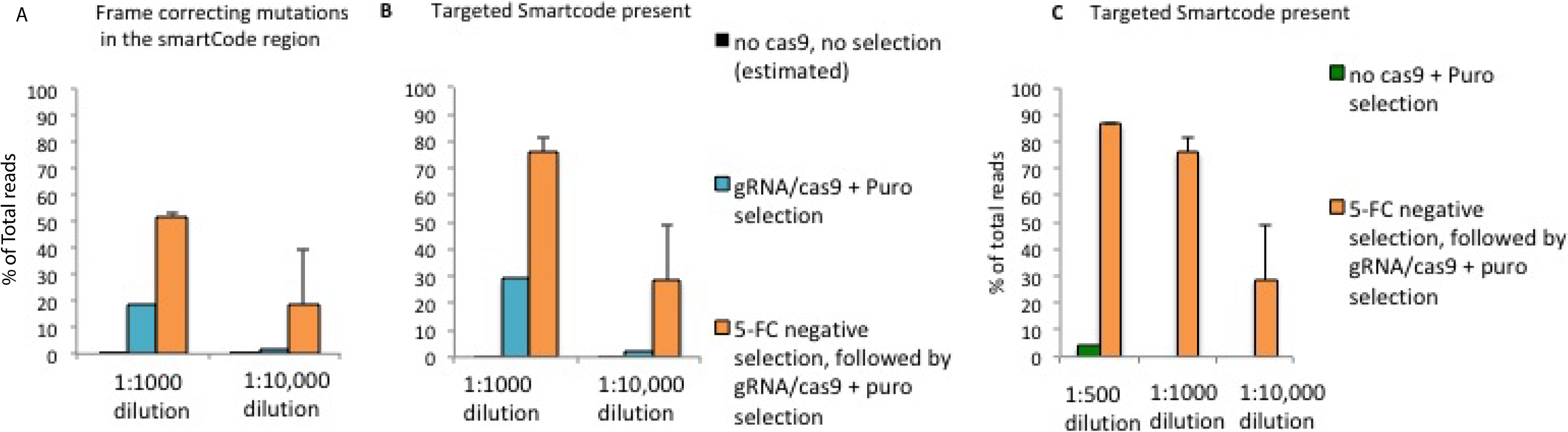
A) and B) Improved enrichment with 5-FC treatment on a population where the majority of cells in the initial population had the GFP SmartCode. Total reads (%) with A) a frame shift mutation in the pasha SmartCode region that will put puromycin in frame or B) with the Pasha SmartCode. C) Enrichment of the target (Pasha) SmartCode through a series of dilutions with the GFP SmartCode in the majority. Error bars show the standard deviation from two replicate experiments.

### A computational approach to SmartCode design

While the GFP and Pasha sites could provide validation of our strategy, we needed to design complex SmartCode libraries that did not target genes that might be present in cells of interest. We therefore began with a highly complex set of synthetic sequences and filtered these against the human and mouse genomes. To select potential SmartCode sequences, we took advantage of an algorithm that we previously developed, CRoatan [13, 14], designed to identify optimal CRISPR target sites in protein-coding genes. As part of the CRoatan workflow, we implemented an analysis of local microhomology to determine the most likely outcome of a repair event [13–15]. We used this approach to identify synthetic sequences that would not only be predicted to be efficiently targeted by Cas9 but also be to generate definable deletions following cas9-mediated cleavage and repair, such that a marker downstream would be shifted from the +2 to the +1 reading frame. We used hamming distance to calculate an orthogonal set of target sites and filtered these sequences for stop codons in the +1 frame, alternative start codons in the +2 frame, and also restriction enzyme sites used in the library cloning process (EcoR1, SalI, NotI, BSIW1). A resulting sequence collection, comprising roughly 50,000 target sites, can be used to create complex SmartCode libraries using in situ oligonucleotide synthesis [16].

### Validation of the SmartCode design strategy

To assess our SmartCode design strategy, we again constructed an experimental simulation. We selected two sequences from our 50K library, termed SmartCode A and SmartCode B and cloned these into a vector that had a ZsGreen marker upstream of the SmartCode and a bicistronic turbo Red Fluorescent Protein (tRFP)-P2A-hygromycin selection marker downstream. This vector was had additional modifications, as described below (Figure 1C).

The reporters were constructed such that the downstream bicistronic cassette was in the +2 reading frame with respect to ZsGreen. A stop codon was included between ZsGreen and the downstream cassette in the +2 frame to prevent read-though from any alternative translational start sites that might be present in ZsGreen. Successful targeting of the SmartCode would move the stop codon out of frame and the selectable markers in to the correct, +1 frame.

Separate Infection of 293T cells with SmartCode A and SmartCode B was followed by FACS purification of zsGreen-positive cells. We combined the two SmartCodes at a 1:1000 ratio, either with SmartCode A in the minority or with SmartCode B in the minority. These mixed populations were then engineered to express an sgRNA targeting the minority SmartCode sequence along with Cas9. After 48 hours, cell populations were selected with hygromycin and surviving cells expanded for a further 3 days. Cells were then FACS purified again for positive expression of zsGreen and tRFP and the resulting SmartCode sequence analysed via Sanger sequencing (Figure 4a-b). Although Sanger sequencing does not allow for determining relative enrichment quantitatively, we observed clear, and interpretable, sequence reads indicating that the majority of the selected population contained the targeted minority barcode. For SmartCode A, we observed, as predicted, a predominant 4 base deletion, and for SmartCode B a predominant 1 base deletion within the SmartCode region.

**Figure 4.**
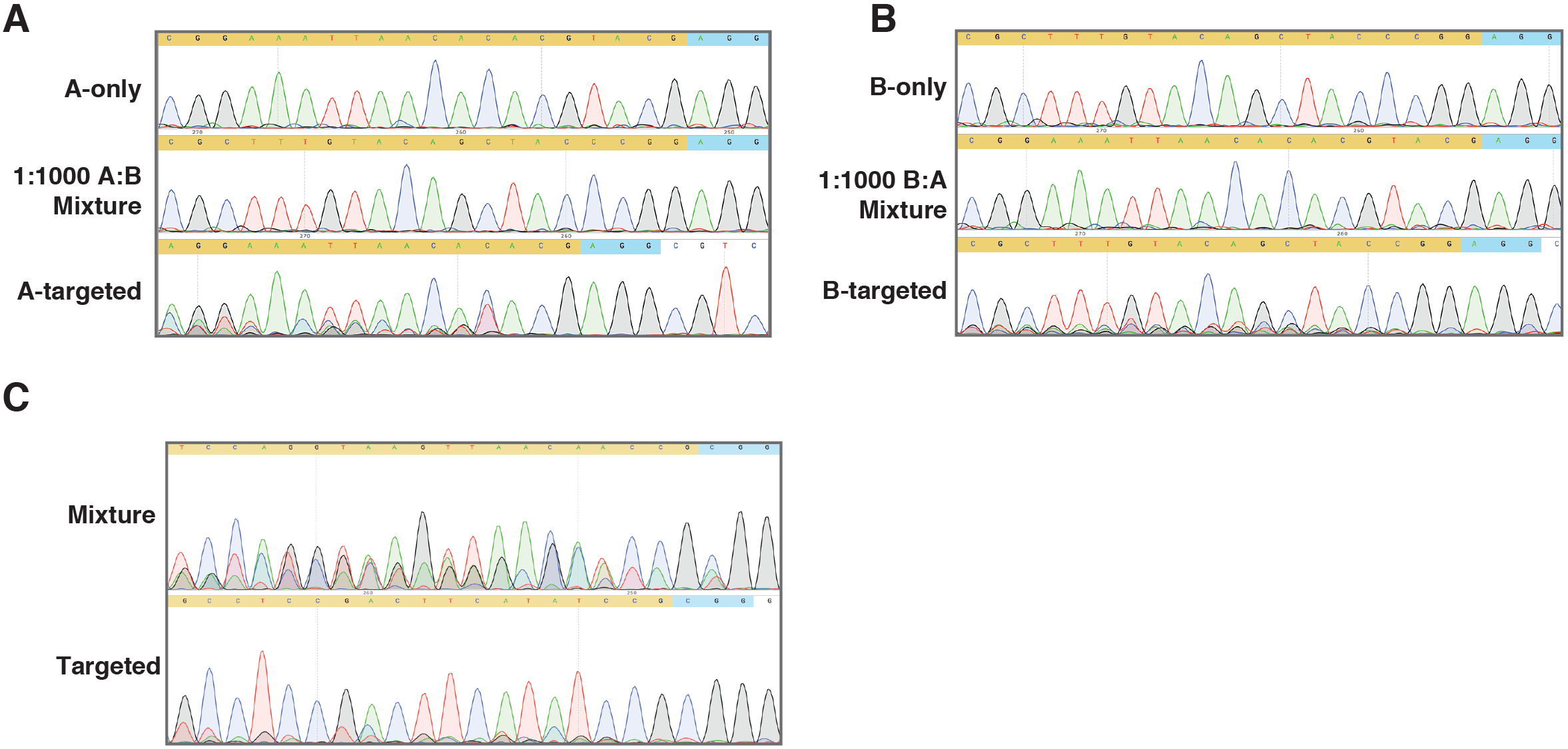
Sanger sequencing reads showing a nonselected cell population containing a majority of Smartcode B in the first instance A) and a majority SmartCode A in the second instance B). Following cas9 retrieval and selection, Sanger reads show the majority of cells contain the (previously in minority) targeted SmartCode. C) Sanger sequencing read from cells containing a complex pool of SmartCodes (~100), followed by cas9 targeting of a single SmartCode.

To determine whether we could isolate a selected SmartCode from a truly complex mixture, we synthesized a library of oligonucleotides corresponding to our computationally predicted sites (as described above) and cloned these into a vector similar to that described above (Fig 1C), and virally infected cells 293T under conditions where each cell would be singly infected. We then simplified this mixture by sorting a population comprising 100 initial founder clones and expanded this for further experiments. We selected one target SmartCode (SmartCode GCCTCCGACTTCATATGCCGCGG) and used a corresponding sgRNA to guide Cas9 for marker activation in that clonal lineage. After applying the dual hygromycin and FACS-based strategy described above, Sanger sequencing of an amplicon comprising the SmartCode region revealed that the majority sequence was derived from the targeted SmartCode and that this contained the predicted −1 deletion (Figure 4C).

### Integrating molecular phenotyping with SmartCode retrieval

A combination of single-cell sequencing and multiplexed, CRISPR-based genetic manipulation has previously been used for high-throughput phenotype determination_[17-19]._In these studies, a conventional barcode sequence was placed adjacent to a polyadenylation signal in the sgRNA expression vector to enable linkage between the targeted gene and the change in transcriptional profile that resulted from its mutation. We took inspiration from the Perturb-SEQ strategy [17] and during the creation of our 50K SmartCode library and coupled a conventional barcode to the SmartCode during synthesis. The set of cloning steps used to create the retroviral library not only appropriately placed the SmartCode within the dual tRFP/Hygro^r^ selection cassette but also sited the second barcode adjacent to a polyadenylation signal, enabling it to be captured in 3′ biased single-cell RNA sequencing workflows.

The same SmartCode library used for the clone retrieval studies described above was used to infect 4T1 cells and, after creating a population founded by 100 initial lineages, these were used in a 10X single-cell sequencing run to produce data on 7000 cells, which following filtering for representation of barcode reads in the singlecell datasets, yielded approximately 700 informative cells. In parallel, PCR amplification was performed to extract a second library to allow the complete set of cell barcode and SmartCode pairings to be verified. 53 founding lineages were found to be represented when using a 10-read cut off. Single-cell data was analysed using the Seurat Package to perform t-distributed stochastic neighbour embedding (t-SNE) clustering. (Fig 5a).

**Figure 5.**
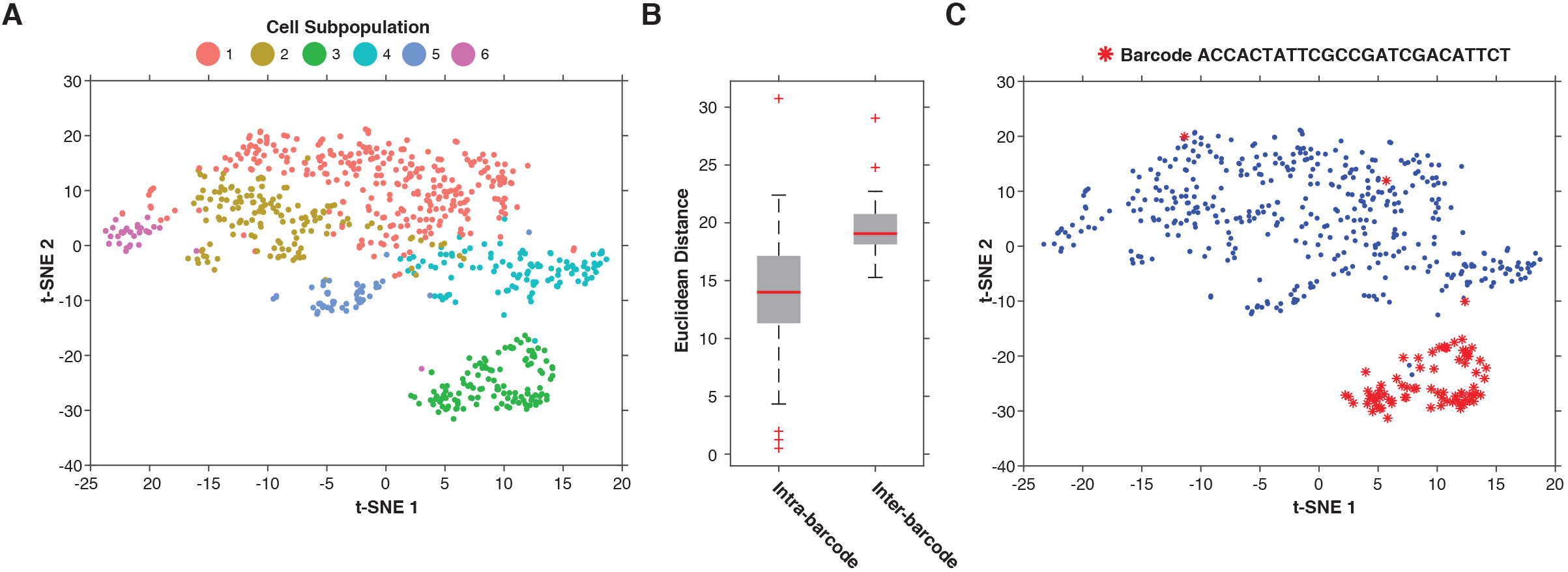
A) T-SNE plot showing clusters of 4T1 cells after single cell 10X sequencing B) Pairwise Euclidean distances between cells using their t-SNE coordinates, showing cells from the same cluster are more likely to have the same linked barcode. C) A distinct population of cells illustrating a common linked barcode.

To determine if these single-cell profiles were capable of distinguishing between different clonal lineages, we calculated all pairwise Euclidean distances between cells using their t-SNE coordinates, and then compared distances between cells containing the same SmartCode versus cells containing different sequences. Indeed, cells harbouring the same SmartCodes showed significantly higher similarity (Figure 5b, p-value = 2.8e^−13^). Furthermore, when the most distinctive cluster in the t-SNE plot was examined, we found that the vast majority of cells within this cluster were derived from the same lineage, as they harboured a single SmartCode (Figure 5c). Based upon our parallel studies (Figure 3 and 4), this lineage could easily be enriched from the starting population using the appropriate guide RNA.

## Concluding remarks

Existing methodologies allow tracking of clonal lineages and their phenotypes in complex and heterogeneous populations. Recent studies have even coupled analysis of clonal populations with genetic perturbations [17]. Here we describe an approach that permits complex populations to be probed for a given behaviour and subsequently permits the retrieval of cells showing that behaviour from within the original mixture. The approach is adaptable to behaviours or phenotypes that are observable at many different levels. Ultimately the incorporation of a genetic barcode that is readable in single-cell RNA sequencing enables not only physical or macroscopic phenotypes to be studied but also molecular phenotypes to be used as a guide for lineage selection.

The general strategy that we describe is extremely flexible. Even in its current incarnation, both positive and negative selections are possible, both in vitro and in vivo. This not only enables the retrieval of clonal lineages from a complex population to facilitate their further study but also the removal of specific lineages from complex mixtures to determine their contribution to the in vitro or in vivo phenotypes of populations, for example, tumour formation, drug resistance, or metastasis.

Though we have validated our approach in a model of limited complexity, we have created robust algorithms that enable complex libraries of SmartCodes to be designed. If one considers heterogeneity in cancer, it is as yet unclear how many clonal lineages are relevant to disease progression. Certainly, studies to date would argue for multiple lineages, perhaps showing genetic or phenotypic diversity [20, 21] that could impact properties relevant to progression and outcome. The methods that we describe perform well beyond the ability to enrich relevant clones from mixtures of the expected relevant complexity, placing the SmartCode strategy in a position to facilitate studies of how cellular heterogeneity impacts nearly every aspect of disease progression.

By further combining SmartCodes with the ability to track and profile lineages by single cell sequencing, we create a powerful suite of technologies for mining the heterogeneity of cell populations in any circumstance where we can individuate cells by barcoding. While obviously applicable to cancer, this approach is also potentially deployable in any number of other biological contexts.

## Materials and methods

### SmartCode Vectors

The SmartCode sequences for Fig 1B were cloned into the MSCV-IRES-GFP (Addgene) retrovirus backbone using primers (1&2) to amplify the puromycin resistance gene with the SmartCode tagged to the forward primer, and cloned with EcoR1 and XhoI. The mutant CodA gene, codAD314A, was cloned using an IDT GBlock and appended to the 3′ end terminus of the puromycin coding sequence using DraIII and XhoI. A stop codon is present at the junction of puromycin and codAD314A when puromycin resistance is out of frame.

For experiments with the SmartCode and linked barcode (Fig 1c) the following sequence scaffold was synthesized:ATCGCCTCCGGCTCCGCCTTGCCCGAATTCSSSSSSSSSSSSSSSSSSSSSGGGTAACGGATCCCTCGACCGGATGTCACCGGTAGATCGTCGCTCGCACGCGTBBBBBBBBBBBBBBBBBBBBBBBBBBGTCTCCTGTGCCTTCTAGTTGCCAGCCATCTGTTG, where the S region contains the SmartCode sequence up to the first base of the PAM and the B region contains the linked barcode for single cell sequencing. Individual constructs were synthesized by IDT, while the complex SmartCode pool was synthesized by CustomArray, Inc. These sequences were amplified with 16 cycles using primers 20 and 21. Amplified sequences were then introduced into a BsmBI linearized backbone backbone via a Gibson reaction. The remaining tRFP-P2A-HygR portion was then ligated via BamHI and MluI into the resulting product, generating the final vector.

### Cell Culture

293T cells and 4T1 cells were maintained in Gibco DMEM supplemented with 10% FBS and 6mls of Pen/Strep. 6ug.ml polybrene was supplemented for each viral infection. 5-FC treatment (90ug/ml) was maintained for 4 days over 1 cell passage. Cells were washed and recovered for 7 days. Cells were passaged for 7 days after Cas9 infection to create mutations. Puromycin selection was done using 2ug/ml over 2 days and cells maintained in this antibiotic supplied media during expansion. Hygromycin selection was done using 200ug/ml for two days, when cells were trypsinized and returned to plate with antibiotic media. Non attached (non cas9 targeted cells) were removed via aspiration.

### Validation of the SmartCode approach

293T cells were infected with the Pasha and GFP SmartCode vectors and FACS sorted from ~20% GFP positive to a purity of 85% GFP positive. A subset of these cells were treated with 5-FC (90ug/ml) for 3 days and left to recover for a further 4 days. A subset of these 5-FC treated cells were then infected with the pLentiCRISPR lentivirus containing an sgRNA for either Pasha (GACAGCGACCATCCGTCCGA) or GFP (GGGCGAGGAGCTGTTCACCG). Cells were maintained for a further 7 days to allow Cas9 to create mutations. A subset of these cells were then treated with puromycin (2ug/ml) and maintained in antibiotic media for a further 7 days to expand the resistant cells. Genomic DNA from treated cells, and corresponding control conditions, was extracted using the PureLink Genomic DNA mini kit (Invitrogen). A ~ 700bp region surrounding the SmartCode was PCR (Primers 3 & 4) amplified in 3 replicate PCR reactions. From this PCR product a ~300bp region was PCR (primers 5 & 6 underlined the Truseq binding sites) amplified to incorporate barcoded NextGen sequencing primers. Each treatment library was PCR barcoded for sequencing with 1 of 12 barcodes (primers 7-19). 12 libraries were pooled per lane of a MiSeq sequencing run and sequence data collected over 101 base pairs.

All virus production was performed as we have described previously [7, 8].

### Validation of the SmartCode design strategy

For experiments with SmartCodes A and B, 293T cells were infected separately with corresponding constructs at a frequency of infection of lower than 10%. Cells were sorted for high zsGreen expression and 3 million cells were used to seed 10cm dishes. After recovery cells were mixed to a ratio of 1:1000 in a total cell count of 9 million and immediately transfected with the target guide RNA (sgRNA) and cas9 (lentiviral construct where the expression of a bicistronic Cas9-blasticidin transcript is driven by the human CMV promoter). The guide RNA was expressed from a human U6 promoter. After continuous culturing for another 72h, the cells were sorted for ZsGreen and tRFP expression to purify the activated cells. This same protocol was also carried out to isolate the single SmartCode from the complex library infected cell mixture; however, here cells were not sorted for RFP expression prior to analyses.

Total genomic DNA was isolated using the DNEasy blood and tissue kit (Qiagen) and the region containing the guide-barcode pairs was amplified using the primers 22 and 23 and purified with a BluePippin (Sage Science), the DNA was analysed by Sanger sequencing using primer 24.

### Miseq analysis

Data was filtered for the presence of shared sequence adjacent to that of the SmartCode region, then the number of bases between these shared sequences was counted and the number of reads for each count were taken. A selection of sequence reads were manually validated for containing the correct deletion in the sequence.

### Single Cell Sequencing Analysis

Single cell sequencing libraries were constructed on the 10X genomics platform, following all suggested procedures and sequenced using the Illumina NextSeq. A single cell count matrix was derived using a pipeline consisting of STAR alignment, UMI tools, and Featurecount.

A second library (primers 25 and 26) was generated from this initial expression library to validate which 10X cell barcodes were associated with which SmartCodes. This library was sequenced on an Illumina MiniSeq platform. Each read pair in this library contained a SmartCode linked barcode, a UMI (representing an initially captured molecule) and a 10x cell barcode. Each 10X cell was assigned a count for each possible SmartCode linked barcode based on the number of unique UMI sequences that were associated with the pair. Cells were filtered based on their being associated with at least 10 SmartCode UMI pairs, where the maximum SmartCode is represented at a ratio of at least 10:1 relative to the next most abundant SmartCode for that cell. SmartCodes were then filtered for those that were associated with at least 10 cells in the 10X data. Cells that were associated with other SmartCodes were removed from subsequent analyses.

After filtering cells based on the quantification of their associated SmartCodes, we applied the Seurat pipeline to cluster the remaining population based on their gene expression profiles. Cells were then labelled in the t-SNE space based on their SmartCode. Euclidean distances were then calculated for all inter-SmartCode cell pairs and also for all intra-SmartCode pairs in this space.

### Primers

1. SCpashapuroF GCG GCG GAA TTC CCA TGG GGA CAG CGA CCA TCC GTC CGA GGG ACC GAG TAC AAG CCC ACG
2. SCgfppuroF GCG GCG GAA TTC CCA TGG GGG GCG AGG AGC TGT TCA CCG GGG ACC GAG TAC AAG CCC ACG
3. puro R xho CGC CGC CTC GAG TCA GGC ACC GGG CTT GCG GGT CAT GCA CC
4. SCGAG F. TCCGCCTCCTCTTCCTCCATCCGC
5. TrueSeq binding primer Forward ACACT CTTTCCCT ACACGACGCCT CTTCCGAT CT CT C CCTTTATCCAGCCCTCACTCCTTCTCTAGGCG
6. TruSeq Binding primer Rev TCGTGACTGAGATTCAGACGTGTGCTCTTCCGATCTACCTTGCCGATGTCGAGCCCGACGCGCGTGAGGA
7. Multiplexing PCR Primer 1.0 AATGATACGGCGACCACCGAGATCTACACTCTTTCC CTACACGACGCTCTTCCGATCT
8. Multiplexing PCR Primer 2.1 CAAGCAGAAGACGGCATACGAGATCGTGATGTGA CTGGAGTTCAGACGTGTGCTCTTCCGATCT
9. Multiplexing PCR Primer 2.2 CAAGCAGAAGACGGCATACGAGATACATCGGTGA CTGGAGTTCAGACGTGTGCTCTTCCGATCT
10. Multiplexing PCR Primer 2.3 CAAGCAGAAGACGGCATACGAGATGCCTAAGTGA CTGGAGTTCAGACGTGTGCTCTTCCGATCT
11. Multiplexing PCR Primer 2.4 CAAGCAGAAGACGGCATACGAGATTGGTCAGTGA CTGGAGTTCAGACGTGTGCTCTTCCGATCT
12. Multiplexing PCR Primer 2.5 CAAGCAGAAGACGGCATACGAGATCACTGTGTGA CTGGAGTTCAGACGTGTGCTCTTCCGATCT
13. Multiplexing PCR Primer 2.6 CAAGCAGAAGACGGCATACGAGATATTGGCGTGA CTGGAGTTCAGACGTGTGCTCTTCCGATCT
14. Multiplexing PCR Primer 2.7 CAAGCAGAAGACGGCATACGAGATGATCTGGTGA CTGGAGTTCAGACGTGTGCTCTTCCGATCT
15. Multiplexing PCR Primer 2.8 CAAGCAGAAGACGGCATACGAGATTCAAGTGTGA CTGGAGTTCAGACGTGTGCTCTTCCGATCT
16. Multiplexing PCR Primer 2.9 CAAGCAGAAGACGGCATACGAGATCTGATCGTGA CTGGAGTTCAGACGTGTGCTCTTCCGATCT
17. Multiplexing PCR Primer 2.10 CAAGCAGAAGACGGCATACGAGATAAGCTAGTGA CTGGAGTTCAGACGTGTGCTCTTCCGATCT
18. Multiplexing PCR Primer 2.11 CAAGCAGAAGACGGCATACGAGATGTAGCCGTGA CTGGAGTTCAGACGTGTGCTCTTCCGATCT
19. Multiplexing PCR Primer 2.12 CAAGCAGAAGACGGCATACGAGATTACAAGGTGA CTGGAGTTCAGACGTGTGCTCTTCCGATCT
20. Amp_FOR ATCGCCTCCGGCTCCGCC
21. Amp_REV CAACAGATGGCTGGCAACTA
22. Genomic_FOR GGCACTTCATCCAGCACAAGCTG
23. Genomic_REV CTACAGCTGCCTTGTAAGTCATTGGTC
24. Sequence CTCGATGAGCTGATGCTTTG.
25. P5_Amp AATGATACGGCGACCACCGAGATCT (binding to the P5 Illumina adapter in the cDNA library)
26. P7_X6_Read2_BC CAAGCAGAAGACGGCATACGAGATTGCACTGTGACTGGAGTTCAGACGTGTGCTCTTCCGATCTGGGCAAAGGAATAGACGCGT (binding on hygromycin, adding Read2, an Index and the other Illumina adapter sequence).

## Acknowledgements

We are grateful to the CRUK CI genomics core for sample sequencing. Support for this project was provided by Cancer Research UK and the DOD Breast Cancer Research Program (grant W81XWH-12-1-0300) and a Winnick Scholar Award.

